# An Optogenetic Toolkit for Light-Inducible Antibiotic Resistance

**DOI:** 10.1101/2022.06.10.495621

**Authors:** Michael B. Sheets, Mary J. Dunlop

## Abstract

Antibiotics are a key control mechanism for synthetic biology and microbiology. Resistance genes are used to select desired cells and regulate bacterial populations, however their use to-date has been largely static. Precise spatiotemporal control of antibiotic resistance could enable a wide variety of applications that require dynamic control of susceptibility and survival. Here, we use light-inducible Cre recombinase to activate expression of drug resistance genes in *Escherichia coli*. We demonstrate light-activated resistance to four antibiotics: carbenicillin, kanamycin, chloramphenicol, and tetracycline. Cells exposed to 465 nm blue light survive in the presence of lethal antibiotic concentrations, while those kept in the dark do not. To optimize resistance induction ranges, we characterize the impact of the promoter, ribosome binding site, and enzyme variant strength using chromosome and plasmid-based constructs. Using time-lapse microscopy, we further show resistance activation dynamics. These optogenetic drug resistance tools pave the way for spatiotemporal control of cell survival.

## Introduction

Antibiotic resistance genes are widely used in synthetic biology. They are included in genetic constructs to ensure plasmid propagation. Resistance genes also play an important role in cloning methods. Examples include chromosomal insertions, where expression of resistance genes can be used as a selective marker for successful integration,^1^ or in the creation of transposon libraries, where drug resistance is used as an intermediate selection mechanism before being swapped for an alternative sequence.^2,3^

Although antibiotic resistance genes are a staple of synthetic biology and microbial biotechnology research, there are few methods for dynamic control of their expression. The ability to control drug resistance spatially and temporally could open new avenues for synthetic biology research. As an analogy, when Sheth *et al*.^4^ developed an inducible origin of replication—another ubiquitous feature within synthetic biology constructs—it sparked new areas of research including biological data storage^5^ and whole-cell riboswitch diagnostics.^6^ Spatiotemporal control over drug resistance could enable dynamic spatial patterning of living biomaterials,^7^ selection of single cells from microfluidic systems,^8,9^ and improved understanding of the role dynamics play in clinical antibiotic resistance.^10^ For example, resistance is often spread through horizontal gene transfer events,^11,12^ which are difficult to monitor and control at the single-cell level. New systems for control offer the potential for future studies quantifying how different spatiotemporal arrangements of cells acquiring resistance can lead to population-level proliferation or collapse.

Optogenetic methods are a powerful and widely used tool for controlling gene expression.^13^ The delivery of light to cells can be regulated in space and time, and can be integrated directly into computational workflows.^14^ Optogenetic systems in bacteria have been used to control gene expression for a variety of applications,^13,15^ including to drive metabolic flux,^16^ regulate the gut microbiome,^17^ and control cell morphology.^18^ Light has also been used to control cell survival with individually designed photo-caged antibiotics^19^ or by leveraging the natural photosensitivity of tetracycline.^20^ However, because these methods require careful protein engineering or exploit properties specific to a single drug, they do not easily generalize across different resistance mechanisms. An alternative approach used a light-inducible promoter to control chloramphenicol resistance.^21^ While such a method could in principle generalize to other resistance genes, experiments were limited to control of the chloramphenicol acetyltransferase enzyme. The ideal platform for light-inducible resistance would be both generalizable for different antibiotic resistance genes and tunable across antibiotic concentrations to flexibly enable diverse studies in synthetic biology and microbiology.

To address these needs, we use the blue light-inducible Cre recombinase OptoCreVvd2 to activate antibiotic resistance genes.^22^ Using this system, we excise a *loxP*-flanked transcription terminator between a gene and promoter, allowing for increased gene expression after exposure to blue light (Fig. 1a). Recombinase technology has been used successfully for a variety of applications that require robust and inducible control of gene expression, including gene logic circuits and cell lineage tracking.^23–25^ We use this system for its relatively short activation time, flexibility in construct design (requiring only the addition of *loxP* sites), and minimal basal expression of activated genes.^22^ In addition, the permanent OFF-ON switch caused by Cre allows for selection of resistant cells at any point after light exposure, allowing for cellular memory after the light input has been removed.

**Figure 1.**
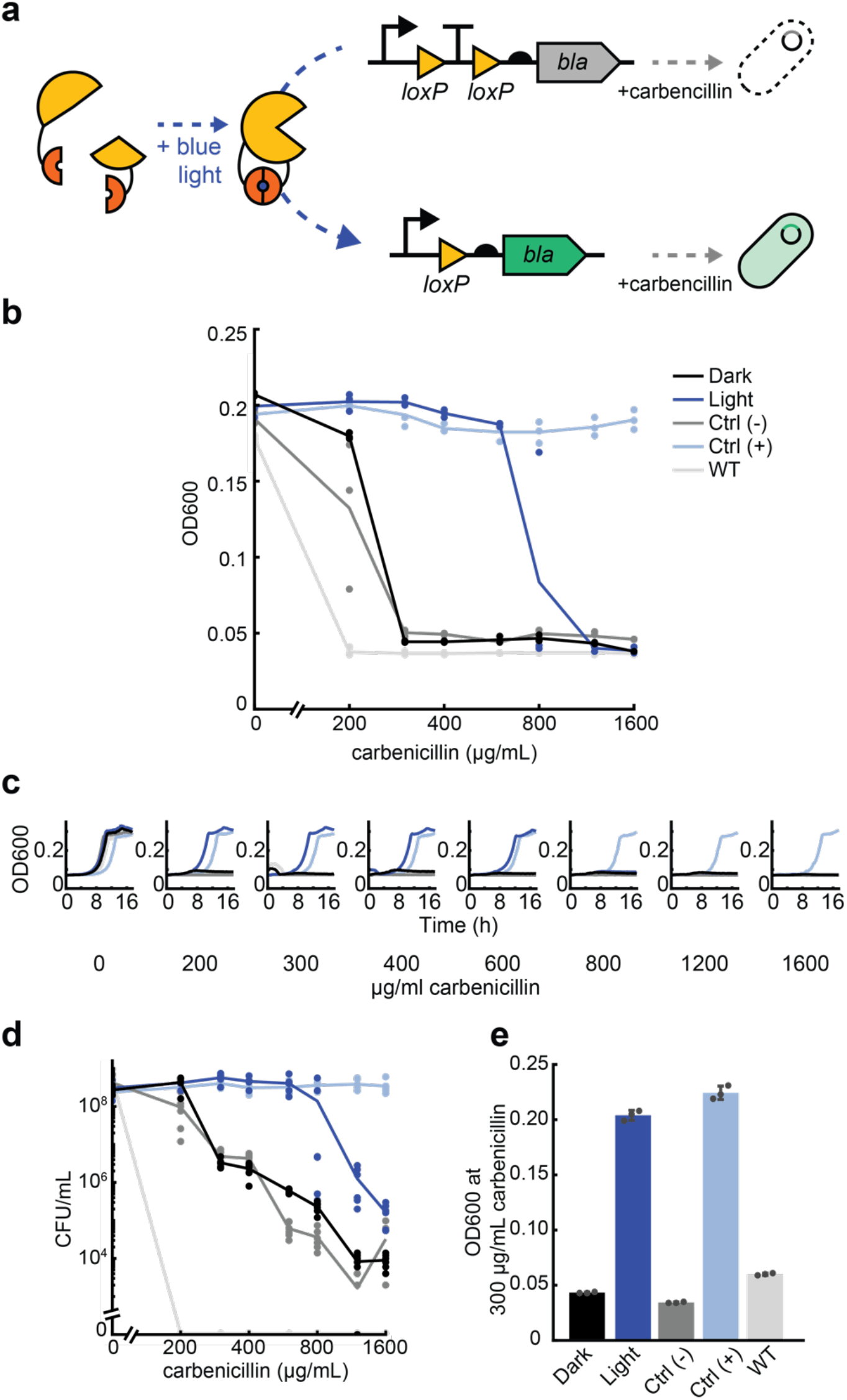
Optogenetic activation of beta-lactamase antibiotic resistance using OptoCre-*bla*. **(a)** Split Cre recombinase fragments are linked to blue light-inducible Vvd photodimer domains. When exposed to blue light, Cre becomes active and can excise a transcription terminator between two *loxP* domains, allowing increased expression of *beta-lactamase* (*bla*). The expression of OptoCre-*bla* then allows cells to survive in the presence of the antibiotic carbenicillin. **(b)** Minimum inhibitory concentration (MIC) curves of chromosomally-integrated OptoCre-*bla* constructs grown in carbenicillin for 18 hours. Light-induced samples were exposed to blue light for two hours immediately before exposure to carbenicillin. Growth measured by OD600 (n = 3). The strains in both the dark and light conditions contain the resistance induction construct and OptoCreVvd2. Control (-) cells contain the resistance activation construct but no Cre recombinase. Control (+) cells contain a constitutively expressed *bla* gene. Wild-type (WT) cells are MG1655 without modification or plasmids. **(c)** Time-course growth of OptoCre-*bla* resistance activation constructs across different concentrations of carbenicillin. **(d)** Colony forming unit (CFU) counts of cultures from MIC data (n = 6). After growth in carbenicillin for 18 hours, samples were spotted on agar plates and colonies were counted the next day. **(e)** Optimal OptoCre-*bla* activation conditions. Resistance activation constructs grow in 300 µg/mL carbenicillin after exposure to blue light for 2 hours, but not when kept in the dark. Growth is quantified by OD600 after 18 hours. Error bars show standard deviation around the mean (n = 3).

Here, we used Cre to induce expression of four antibiotic resistance genes, which we selected for their ubiquity in synthetic biology applications as well as their range of mechanisms of action (Table 1). Specifically, we chose the carbenicillin/ampicillin resistance gene beta-lactamase (*bla*), which is both clinically relevant and widely used in synthetic biology. Beta-lactam antibiotics inhibit cell wall biosynthesis, and are enzymatically degraded by the beta-lactamase enzyme.^12,26,27^ We also selected kanamycin nucleotidyltransferase (*knt*), which provides enzymatic resistance against kanamycin, an antibiotic that causes mistranslation by the 30S ribosomal subunit.^28^ Chloramphenicol acetyltransferase (*cat*) provides enzymatic resistance against chloramphenicol, which interferes with the 50S ribosomal subunit to cause protein synthesis to stall.^29^ Lastly, we included the tetracycline efflux pump A (*tetA*) as a non-enzymatic, efflux-based resistance mechanism.^30^ These four antibiotics include both bactericidal (carbenicillin/ampicillin and kanamycin) and bacteriostatic (chloramphenicol and tetracycline) drugs. This selection of resistance mechanisms shows both the broad variety of mechanisms that can be controlled using this system and introduces multiple options for synthetic tools that can be incorporated into existing bacterial systems.

**Table 1.**
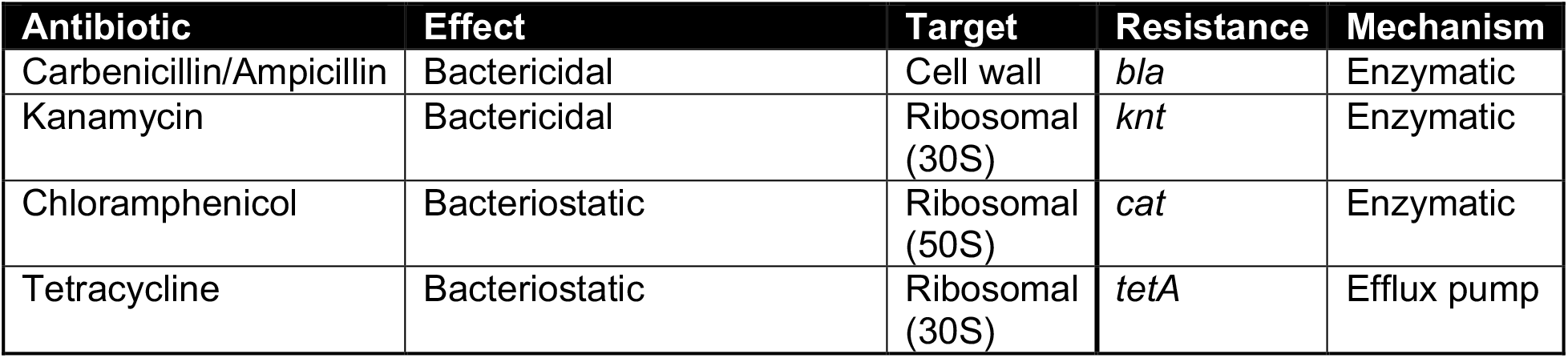
Summary of antibiotics and corresponding resistance genes used in this work.

**Table 2.** Resistance activation constructs used in this study and respective dark and light state MICs. Optimal constructs for each antibiotic are highlighted in gray, where this selection considers both fold change and total numerical difference in MIC on light exposure.

| Resistance | Site | Promoter | RBS | Dark MIC (µg/mL) | Light MIC (µg/mL) |
| --- | --- | --- | --- | --- | --- |
| **OptoCre-*bla*** |  |  |  |  |  |
|  | chromosomal (*nupG*) | *P* | *R* | 300 | 1200 |
|  | plasmid (p15A origin) | *P* | *R* | 10000 | 10000 |
| **OptoCre-*knt*** |  |  |  |  |  |
|  | chromosomal (*nupG*) | *P\*\** | *R* | 200 | 600 |
|  | chromosomal (*nupG*) | *P\** | *R* | 600 | 1200 |
|  | chromosomal (*nupG*) | *P* | *R* | 1600 | 2400 |
|  | chromosomal (*nupG*) | *P* | *R\** | 4 | 6 |
|  | plasmid (p15A origin) | *P\*\** | *R* | 4800 | 6400 |
|  | plasmid (p15A origin) | *P\** | *R* | 4800 | 6400 |
|  | plasmid (p15A origin) | *P* | *R* | 6400 | 9600 |
|  | plasmid (p15A origin) | *P* | *R\** | 15 | 40 |
| **OptoCre-*cat*** |  |  |  |  |  |
|  | chromosomal (*nupG*)<br>*cat* | *P* | *R* | 15 | 30 |
|  | plasmid (p15A origin)<br>*cat* | *P* | *R* | 150 | 300 |
|  | chromosomal (*nupG*)<br>*cat*<sub>T172A</sub> | *P* | *R* | 10 | 30 |
|  | plasmid (p15A origin)<br>*cat*<sub>T172A</sub> | *P* | *R* | 75 | 200 |
| **OptoCre-*tetA*** |  |  |  |  |  |
|  | chromosomal (*nupG*) | *P* | *R* | 1.5 | 1.5 |
|  | plasmid (p15A origin) | *P* | *R* | 1.5 | 1.5 |
|  | chromosomal (*nupG*) | *P<sub>tet</sub>* | *R<sub>tet</sub>* | 1 | 2 |
|  | plasmid (p15A origin) | *P<sub>tet</sub>* | *R<sub>tet</sub>* | 4 | 8 |

When controlling expression of antibiotic resistance genes, key performance metrics include the uninduced and induced expression levels. For example, when using potent enzymes like beta-lactamases, even a small amount of basal expression can allow bacterial growth in the presence of an antibiotic. Moreover, expression in the induced state should also be sufficient to provide resistance at drug concentrations comparable to typical working ranges for the antibiotic, which could further vary between use cases. To optimize these two features in our platform, we varied the gene copy number, promoter, ribosome binding site (RBS), and coding sequence to tune the minimum inhibitory concentration (MIC) of antibiotic at which cells survive after exposure to blue light, while maintaining basal expression levels that are low enough to avoid erroneously triggering survival. We further demonstrated live activation of resistance genes and characterized cellular responses using single-cell time-lapse microscopy. This work builds on the long use of antibiotics as cellular control mechanisms in synthetic biology, adding a spatial and temporal control mechanism to existing systems, setting the stage for future applications where light is used in combination with antibiotics to enable flexible control of cell behavior and survival.

## Results

For optogenetic control of resistance genes, we used the blue light-inducible split Cre recombinase OptoCreVvd2.^22^ This system allows for excision of genetic elements placed between *loxP* sites when cells are exposed to blue light. Excision can be completed in approximately two hours, which is comparable to or faster than many existing bacterial optogenetic systems.^18,21,31,32^ We used OptoCreVvd2 to excise a transcription terminator placed inside *loxP* sites between a promoter and an antibiotic resistance gene, allowing for expression of the resistance gene only after exposure to blue light.

We first used this system to control transcription of the beta-lactamase (*bla*) resistance gene (which we denote ‘OptoCre-*bla’*, Fig. 1a). Beta-lactam antibiotics, including ampicillin and carbenicillin, inhibit peptidoglycan layer biosynthesis in the bacterial cell wall. Beta-lactamase enzymes can inactivate beta-lactam antibiotics by hydrolyzing the beta-lactam ring on the antibiotic.^33^ For these studies, we used the TEM-116 beta-lactamase, which is commonly used in antibiotic resistance cassettes for plasmid selection.^34^ We integrated this genetic construct after *nupG* in the *E. coli* MG1655 chromosome.

To measure light-induced antibiotic resistance, we exposed cultures of OptoCre-*bla* to blue light for two hours, then grew them overnight in the presence of carbenicillin and compared growth to cultures kept in the dark. We observed blue light-dependent differences in cell proliferation, where the minimal inhibitory concentration (MIC) necessary to prevent growth was 300 µg/mL for cultures kept in the dark and 1200 µg/mL for cultures exposed to blue light (Fig. 1b). Negative control (- Control) cells with only the reporter and no Cre recombinase had nearly identical survival to cells with the full construct grown in the dark, indicating low expression of *bla* in the uninduced state. Positive control (+ Control) cells with constitutive expression of *bla* from its native promoter and RBS grew in all concentrations of carbenicillin, as expected for a fully resistant strain. We further included *E. coli* MG1655 as a wild-type negative control, which did not grow in any concentration of carbenicillin used here.

To confirm that blue light-induced cells grow at rates comparable to the positive control strain, we collected time-series data demonstrating normal growth rates under a broad range of carbenicillin concentrations for cells grown in blue light, while cells without light induction failed to grow (Fig. 1c). We further validated the optical density-based MIC data by using colony forming unit (CFU) counts after antibiotic exposure (Fig. 1d). Although MIC data is an accurate assessment of cell growth in the presence of antibiotic, beta-lactam antibiotics also cause cell filamentation, which can increase optical density (OD) readings even when cells are not dividing, making *bla* resistance specifically important to confirm by CFU measurement.^35^ However, our results using CFU counts confirm that the optical density measurements also translate to a clear difference in cell survival.^36^ Overall, we found that OptoCreVvd2 can be used to precisely induce beta-lactamase resistance and identified concentrations of carbenicillin with robust differences between dark and blue-light activated expression of the *bla* resistance gene construct (Fig. 1e).

We next sought to generalize this system to other antibiotic resistance genes. While the excision of a terminator can easily couple light to the expression of an antibiotic resistance gene, this is not the same as light-inducible survival. Control of survival requires no or low expression of the antibiotic resistance gene in the dark, such that cells remain susceptible to antibiotics. The design also requires that induction of the resistance gene is sufficient to confer resistance. The thresholds for these two features can vary dramatically with different antibiotic resistance mechanisms, which have different rates of antibiotic degradation and export. Thus, while previous work in our lab has shown that OptoCreVvd2 can control expression of a fluorescent protein with low basal expression and 10-fold change with light,^22^ naively replacing the fluorescence gene with an antibiotic resistance gene may not produce the desired behavior. Therefore, we defined a general process for adapting the OptoCreVvd2 system to different antibiotic resistance genes (*bla, knt, cat*, and *tetA*) and were able to show survival at customized ranges of antibiotic concentration.

To generalize our system to other antibiotics, we began by exchanging *bla* for *kanamycin nucleotidyltransferase* (*knt*) in our induction construct to make OptoCre-*knt* (Fig. 2a). The antibiotic kanamycin causes mistranslation by binding to the 30S subunit of the bacterial ribosome. The *knt* enzyme catalyzes transfer of a nucleotide to kanamycin, inactivating the antibiotic.^28^ Although our initial design of OptoCre-*knt* did show resistance activation using light, the kanamycin concentration required to see this difference was very high – over 1000 µg/mL (Fig. 2b), compared to 25-50 μg/mL commonly used for plasmid propagation.^20,34^ Although matching common working concentrations of antibiotics is not essential, using concentrations in the vicinity of these ranges provides benefits including the ability to study community effects at physiological concentrations and limiting overall antibiotic needs.

**Figure 2.**
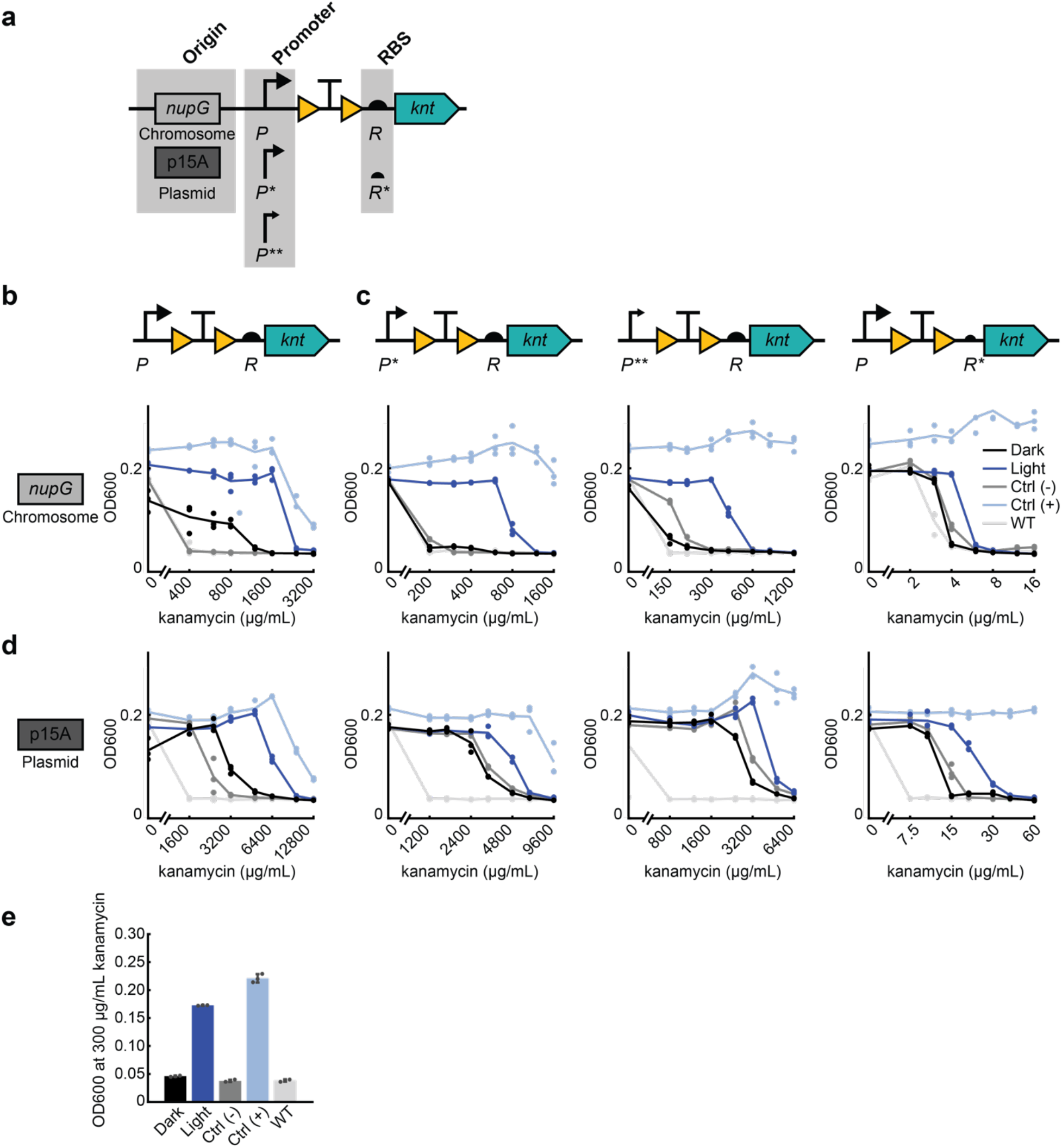
Optimizing optogenetic activation of kanamycin resistance from OptoCre-*knt* by modifying promoter and ribosome binding site (RBS). **(a)** Expression levels of the kanamycin resistance gene *knt* can be tuned by changing the strength of the promoter and RBS, as well as the origin of replication. Promoter strength ranges from P (medium) to P* (medium-low) to P** (lwo). RBS strength ranges from R (strong) to R* (weak). **(b)** MIC curves of OptoCre-*knt* activation cassettes on the chromosome, with promoter P and RBS R (n = 3). **(c)** MIC curves of OptoCre-*knt* activation cassettes on chromosome, with weaker promoters P* or P**, or weaker RBS R*. **(d)** MIC curves of OptoCre-*knt* activation cassettes on a plasmid with the p15A origin of replication. **(e)** Optimal OptoCre-*knt* activation conditions, using P* and R on the chromosome at 300 µg/mL kanamycin. Growth is quantified by OD600 after 18 hours. Error bars show standard deviation around the mean (n = 3).

Thus, we set out to tune gene expression through optimization of the genetic architecture surrounding the gene. To lower basal expression, we weakened the promoter or RBS driving *knt* expression. First, by changing the promoter and RBS of OptoCre-*knt*, we were able to shift the expression of the resistance gene to allow survival at antibiotic concentrations much closer to the MIC of wild-type MG1655 (Fig. 2c). We tested a range of promoter and RBS combinations to show how these alterations impact survival at varying antibiotic concentrations. We used constitutive promoters of varying strength all based on the T7A1 viral promoter, varying from medium, P, to medium-low, P*, to low, P**, transcriptional strength. We also used the RBS of gene 10 in the T7 phage,^37^ which we denote R, as well as a RBS that we computationally designed to be weaker,^38^ which we denote R* (Fig. 2a). Changing P to P* decreased the MIC for the dark state to 200 µg/mL kanamycin, while P** further reduced it to 150 µg/mL (Fig. 2c). The MIC for the light state was also reduced, as expected, but still maintains a wide range of kanamycin concentrations resulting in survival. With P*, kanamycin levels between 200 and 800 µg/mL result in light-induced survival, while for P** the range is from 150 to 500 µg/mL. Using R* in combination with P caused a dramatic decrease in both the dark state MIC and the effective concentrations for light-induced survival, resulting in a narrow range between 4 and 6 µg/mL kanamycin.

Although the chromosomally integrated constructs used so far have the advantages of low background expression and do not require a selection marker, plasmids offer their own advantages for light-inducible resistance systems. Many resistance genes are naturally found on plasmids, and a plasmid origin allows for convenient transfer of systems between different strains. We further characterized our constructs on plasmids containing the p15A origin of replication, which has approximately ten copies per cell (Fig. 2d).^39^ Changing from chromosomal integration to a p15A plasmid increased the range of antibiotic concentrations at which cells containing OptoCre-*knt* selectively survive by over 5-fold. Despite this increase, strategies such as lowering promoter or RBS strength can have a counterbalancing effect. We also characterized a p15A plasmid-based OptoCre-*bla*, however its basal resistance was too high to be considered functional (Fig. S1a). Overall, the added functionality provided by a plasmid-based system with OptoCre-*knt* increases the ease at which these constructs can be used in different strains or contexts.

The flexibility afforded by these different designs led us to develop multiple constructs, and the optimal construct is likely to be application-specific. For example, the lower concentrations shown here are near the wild-type MIC, which is optimal for studies looking to characterize resistance acquisition using phenotypically-relevant antibiotic concentrations. In contrast, the higher concentrations allow more stringent cell selection for studies where only the activated cells should survive. Through this optimization process, we found constructs that allow a greater kanamycin MIC fold change between dark and light-exposed cultures compared to our original construct, notably P* and R driving OptoCre-*knt* expression on the chromosome is an ideal example of our optimized design (Fig. 2e).

Light-induced survival requires a tight OFF-state where cells are susceptible to antibiotic. As we have demonstrated, this can be achieved with low basal expression of the resistance gene. However, the uninduced state can also be minimized if the resistance enzyme itself is weaker. Thus, when activating the chloramphenicol acetyltransferase (*cat*) enzyme, we took advantage of a known mutation to decrease the strength of the enzyme itself. Chloramphenicol prevents protein synthesis by binding to the 50S ribosomal subunit, where it inhibits peptide bond formation. The *cat* enzyme prevents chloramphenicol from binding to the ribosome by attaching an acetyl group from acetyl-CoA to the antibiotic.^40^ By using the weaker *cat*_*T172A*_ variant,^41,42^ we lowered the concentration of antibiotic at which cells survive (Fig. 3a). In the OptoCre-*cat* design, we used the promoter P and RBS R, and compared light-induced survival by *cat* and *cat*_*T172A*_. Here we observed a sharp decrease in dark OFF-state resistance with *cat*_*T172A*_ compared to *cat*, lowering basal resistance to that of the wild-type strain on a chromosomally integrated construct (Fig. 3b-c). We also observed a decrease in MIC values for *cat*_*T172A*_ on a p15A plasmid origin (Fig. 3d). This enzyme mutant approach to optimization may be particularly helpful when working with enzymes that show resistance to high concentrations of antibiotic even with minimal basal gene expression, or if using this system on a plasmid with a high copy number where it is hard to limit basal expression. This approach creates another point at which resistance levels can be fine-tuned, and the light-induced growth difference shown by *cat*_*T172A*_ on a p15A plasmid origin is a particularly versatile optimized design (Fig. 3e).

**Figure 3.**
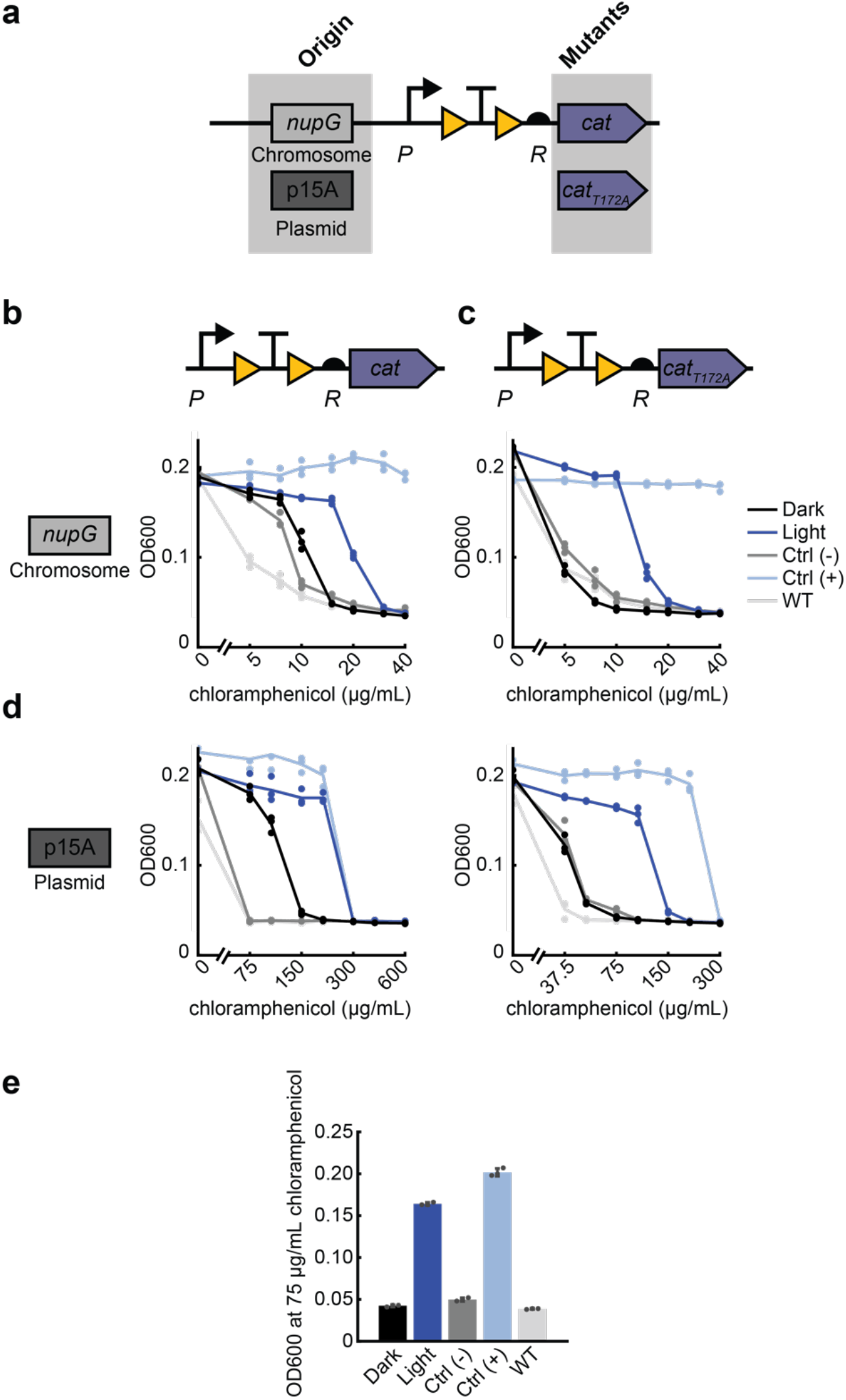
Tuning optogenetic activation of chloramphenicol resistance from OptoCre-*cat* through decreased enzymatic activity. **(a)** Resistance given by the chloramphenicol resistance gene *cat* can be lowered by using the T172A variant on a chromosomal or plasmid origin. **(b)** MIC curves of OptoCre-*cat* activation cassette on the chromosome, with the native enzyme (n = 3). **(c)** MIC curves of *cat*_*T172A*_ activation cassette on the chromosome. **(d)** MIC of the *cat* and *cat*_*T172A*_ activation cassettes on a plasmid with the p15A origin. **(e)** Optimal activation conditions, using *cat*_*T172A*_ with promoter P and RBS R on a plasmid origin at 75 µg/mL chloramphenicol. Growth is quantified by OD600 after 18 hours. Error bars show standard deviation around the mean (n = 3).

Light induction can also be applied to non-enzymatic antibiotic resistance mechanisms, such as the *tetA* efflux pump. The antibiotic tetracycline reversibly binds to the 30S ribosomal subunit, inhibiting protein synthesis. The *tetA* efflux pump localizes to the inner membrane and exports magnesium-tetracycline chelate complexes by importing a proton.^43^ In sharp contrast to *bla, knt*, and *cat* where the native resistance levels were high and our engineering efforts aimed at reducing potency, we found that our initial design for inducible *tetA* did not show resistance over wild-type MG1655 when expressed with promoter P and RBS R on a p15A origin plasmid (Fig. S1b), conditions which produced the highest levels of resistance in constructs tested previously. To compensate for this, we opted to use the strong native promoter P_tet_ with its corresponding RBS R_tet_ to allow full expression of the *tetA* gene to create OptoCre-*tetA* (Fig. 4a).^44^ Using this native architecture, OptoCre-*tetA* showed some activation when chromosomally integrated (Fig. 4b), and exhibited strong activation on the p15A plasmid origin (Fig. 4c). Notably, for both designs, the dark-state basal resistance over wild-type MG1655 was very minimal, allowing for activation of OptoCre-*tetA* at low tetracycline concentrations. Thus, we found that the p15A plasmid-based version is an ideal construct for tetracycline resistance (Fig. 4d).

**Figure 4.**
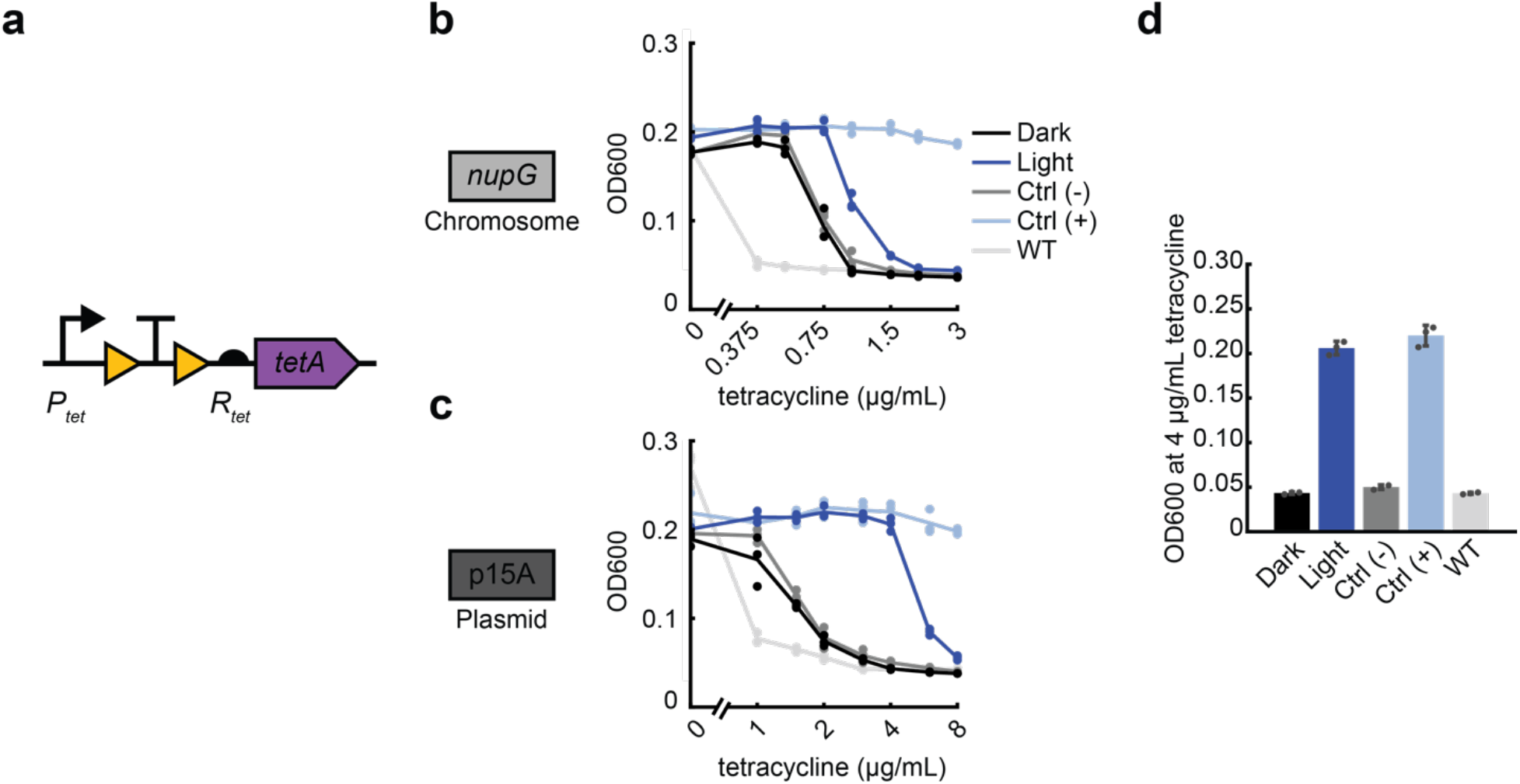
Optogenetic activation of efflux-based tetracycline resistance with OptoCre-*tetA*. **(a)** Tetracycline resistance gene *tetA* is expressed using the native promoter P_tet_ and native RBS R_tet_, to allow maximal expression of the gene. **(b)** MIC of the OptoCre-*tetA* activation on the chromosome and **(c)** p15A plasmid origin (n = 3). **(d)** Optimal OptoCre-*tetA* activation conditions, using the p15A plasmid origin at 4 µg/mL tetracycline. Growth is quantified by OD600 after 18 hours. Error bars show standard deviation around the mean (n = 3).

The spatial and temporal precision enabled by optogenetics allows these constructs to be used for a variety of applications, including single-cell studies of bacterial antibiotic resistance. How resistance acquisition leads to bacterial survival at the single-cell level is of particular interest in the context of horizontal gene transfer. Existing single-cell horizontal gene transfer studies have previously characterized gene transfer rates and shown important connections to quorum sensing.^45,46^ However, natural instances of horizontal gene transfer are infrequent and difficult to control in space and time, especially relative to antibiotic exposure. Previous studies have also shown that stochastic acquisition of resistance in single cells is not always enough to cause the proliferation of phenotypically resistant cells.^47^ To study when horizontal gene transfer events lead to the spread of resistance in populations, it would be interesting to model a single cell’s acquisition of resistance with optogenetic control of antibiotic susceptibility. This could be used to characterize when and how resistance acquisition in single cells leads to antibiotic evasion, and how specific antibiotic dosing schedules and concentrations impact evasion frequency.

Here we show a proof-of-concept for the first steps in this class of studies by inducing cell growth using blue light for cells on agarose pads containing antibiotic. Using time-lapse microscopy, we placed cells containing light-activatable antibiotic resistance on agarose pads containing antibiotics and compared the growth of cells that were kept in the dark to those exposed to blue light. For these studies we elected to focus on a subset of resistance genes, selecting kanamycin and chloramphenicol as examples of bactericidal and bacteriostatic antibiotics, respectively. We characterized resistance activation for chromosomally integrated OptoCre-*knt* resistance to kanamycin (Fig. 5a, Movie S1), and plasmid-based OptoCre-*cat* resistance to chloramphenicol (Fig. 5b, Movie S2). We found that cells with light-induced resistance show a short lag before growth compared to their constitutively resistant positive controls, which likely corresponds to the time needed to excise the transcription terminator and allow expression of the resistance gene. In contrast, when cells were kept in the dark, we observed no growth and examples of loss of membrane integrity (Movie S1-2). The use of microscopy and analysis here shows how populations of cells that dynamically acquire resistance can survive antibiotic exposure. This work shows the potential of using light to control antibiotic resistance, and in the future could be modulated temporally to show how the timing of resistance acquisition impacts survival, complementing studies such as work by Koganezawa *et al*.^10^ that have revealed a non-trivial mechanism of physiological adaptation to resistance gene loss driven by ribosomal subunit balancing. Alternatively, light can be applied spatially to show how user-specified geometric arrangements of resistant cells affect growth of their susceptible neighbors.

**Figure 5.**
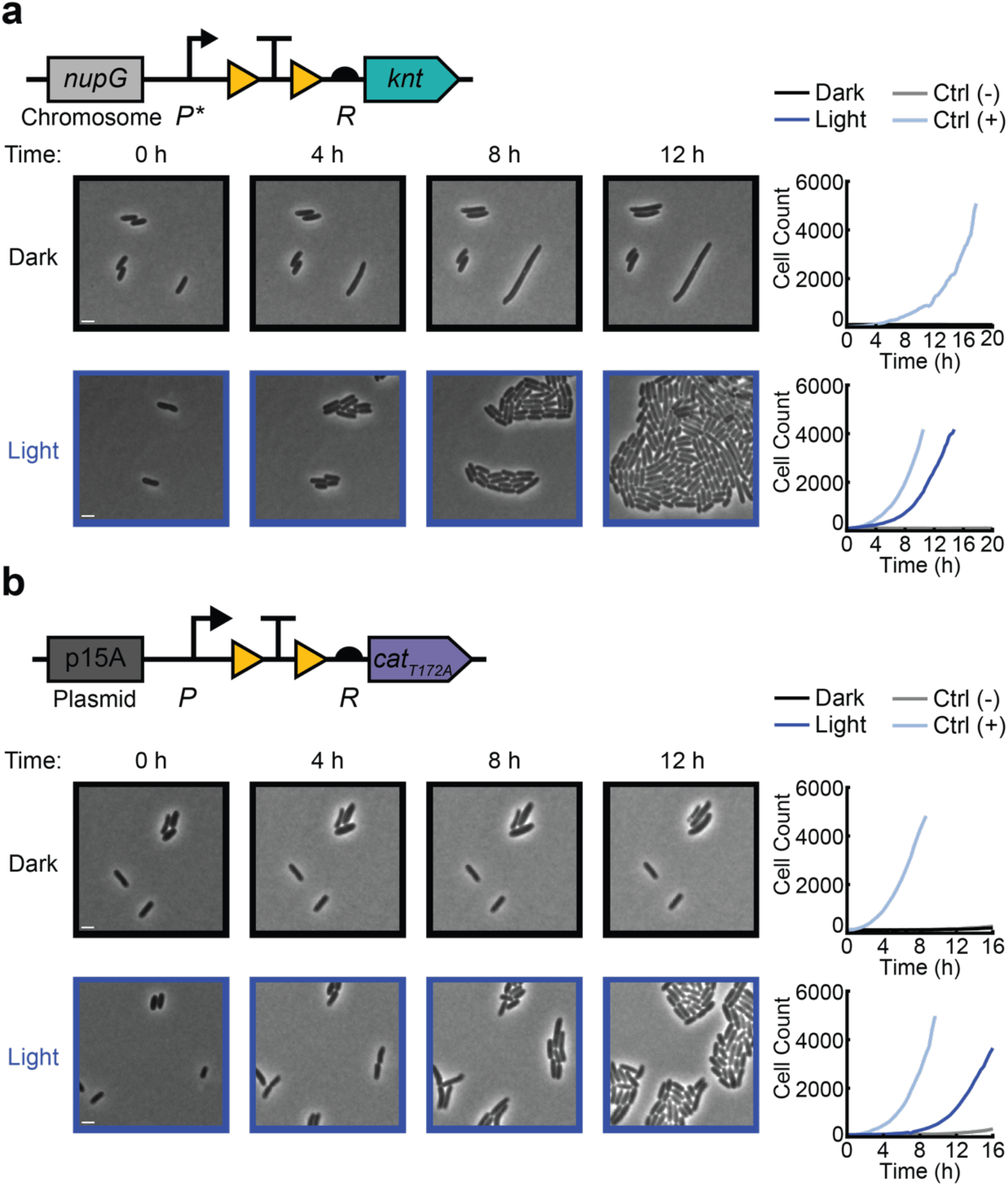
Single-cell time-lapse microscopy of light-induced antibiotic resistance. Activation of **(a)** the chromosomal OptoCre-*knt* resistance construct with promoter P* and RBS R on agarose pads containing 400 µg/mL kanamycin, and **(b)** the p15A plasmid-based OptoCre-*cat*_*T172A*_ resistance construct with promoter P and RBS R on agarose pads containing 60 µg/mL chloramphenicol. Microscopy images show representative samples of the resistance activation strains without or with blue light (scale bar = 2 µm). Cell counts over time are cumulative across multiple imaging positions for each condition, with each plot containing the OptoCre resistance strain along with negative and positive controls (n = 3).

## Discussion

We have developed and optimized optogenetically controlled systems for four antibiotic resistance genes. Using a blue light-inducible Cre recombinase, we have shown activation of *bla, knt, cat*, and *tetA* to induce resistance over a range of antibiotic concentrations. These resistance genes span multiple mechanisms and represent antibiotics and resistance genes commonly used in synthetic biology and microbiology labs. A crucial aspect of designing inducible resistance is the level of expression in the uninduced and induced states, whose optimal levels vary dramatically with the strength of the resistance gene. Therefore, we tested promoters, RBSs, and enzyme mutants of varying strength to find constructs that show optimal basal expression and fold changes of resistance activation. To control copy number at the DNA level, we also compared constructs chromosomally-integrated in the *nupG* region and on a p15A medium copy plasmid, which improved flexibility in experimental design. We found that optimizing resistance induction constructs at the promoter, RBS, enzyme, and copy number levels can be used as a generalizable approach, providing multiple options for altering expression to allow for flexibility in construct design to meet experimental contraints. For example, studies which require a native promoter, specific plasmid origin, or specific resistance protein, can be accommodated as this system is adapted to specific use cases or other resistance genes.

We selected the four antibiotic resistance mechanisms here to be broadly applicable for both synthetic biology and microbiology uses. These antibiotics and resistance genes are often used as plasmid selection markers and in synthetic biology control systems.^20,34^ As an example of an application, these constructs have the potential to be applied to single-cell selection in microfluidic systems. Selection of single cells of interest from a microfluidic device is a challenge, and current methods require complex optical traps and valve-based microfluidic devices.^9^ Combining light-inducible resistance with a digital micromirror device for precise targeting of light would allow for antibiotic selection of a cell line of interest from device outflow, without any chip modification or additional hardware requirements beyond light exposure. Using light also allows integration into computational workflows, potentially facilitating automated screening and selection for phenotypes of interest, where selected cells retain resistance expression and can be harvested for characterization at any point after activation. Additionally, given the prevalence of recombinases in cellular logic,^24^ this system could be modularly integrated with larger recombinase circuits as a logic output controlling cell survival. For instance, recombinase-controlled resistance genes could be broadly used to regulate cell survival only if certain environmental conditions (small molecule, light, or temperature) are met.^24^ As both recombinases and the selected resistance genes are commonly used in synthetic biology, this allows for optogenetic control to be straightforwardly incorporated into existing genetic systems.

Further, we envision these light-inducible antibiotic resistance genes being useful as a synthetic system for studying horizontal gene transfer. These systems are well-suited to examine how single-cell resistance gene acquisition events lead to population expansion or decline. Being able to reliably activate resistance gene “acquisition” using light removes the need to rely on observations of infrequent and stochastic natural horizontal gene transfer events. Notably, *bla* is particularly problematic in clinical settings,^27^ and known to be transferred extensively when the gut is exposed to ampicillin.^12^ Recent studies looking at single-cell instances of horizontal gene transfer have also shown that quorum sensing and biofilm structures can be important in initiating transfer events,^45,46^ however experiments thus far have been limited to the study of infrequent, stochastic horizontal transfer events. Critically, because not all resistance acquisition events lead to the proliferation of resistant populations,^47^ synthetic control of gene transfer could reveal when and how cells acquiring resistance proliferate. This level of spatial and temporal control could allow researchers to determine what cellular arrangements and antibiotic treatment schedules lead to expansion or collapse of microbial communities consisting of resistant cells and their susceptible neighbors.^47,48^

Antibiotic resistance genes are ubiquitous and fundamental tools of synthetic biology. However, there are few current methods that allow dynamic regulation of their expression. The ability to induce resistance using light at the single-cell level expands this core tool, improving spatiotemporal control and enabling new applications in cellular selection and the study of antibiotic resistance acquisition.

## Methods

### Strains and plasmids

All antibiotic resistance assays use *E. coli* strain MG1655. We constructed plasmids using the Gibson assembly method,^49^ or using Golden Gate assembly^50^ in cases where the constructs contained fragments under 100 base pairs in length (Table S1).

Chromosomally integrated constructs were inserted using the Lambda Red recombinase system^1^ downstream of *nupG* (forward homology site: GGTTCTGGCCTTCGCGTTCATGGCGATGTTCAAATATAAACACGT, reverse: GGCGTGAAACGGTTGTACGGTTATGTGTTGAAGTAAGAATAA). Antibiotic resistance cassettes flanked with FRT sites were used to select successful integrations (we used *knt* for *bla* and *cat* activation constructs, *cat* for *knt* and *tetA* activation constructs); these cassettes were then cured using a plasmid-based FLP recombinase on a temperature sensitive origin following the protocol from Datsenko and Wanner.^1^ Finally, we cured the temperature sensitive plasmid before subsequent experiments.

The plasmids used for OptoCreVvd2 expression are derived from Sheets *et al*.^51^ For *bla* activation studies, we changed the plasmid selection cassette from *bla* to *cat*, using the gene from the BglBrick plasmids^34^ and primers listed in Table S1. The antibiotic resistance activation plasmids were made by changing the respective promoter, RBS, and reporter gene of pBbAk-W4-loxTTlox-mRFP1 from Sheets *et al*.^22^ Plasmid origins of replication and sequences for *bla, knt*, and *cat* were taken from the BglBrick plasmid series.^34^ The sequence for *tetA* was obtained from AddGene plasmid #74110 (pRGD-TcR) deposited by Hans-Martin Fischer.^52^ All plasmid-based resistance constructs contain the p15A origin. Plasmid-based constructs for *knt* activation contain *cat* as a selection marker, and constructs for *bla, cat*, and *tetA* activation contain *knt* as a plasmid selection marker. Positive controls for *bla* and *tetA* contain the respective gene with its native promoter and RBS on a plasmid with the p15A origin. Positive controls for *knt* and *cat* contain the respective gene with its native promoter and RBS on a plasmid with the p15A origin for studies using a plasmid-based reporter, or integrated into the *nupG* region for studies using chromosomally integrated reporters. Promoters used were medium, medium-low, and low-strength variants of T7 A1,^53^ denoted P (TTATCAAAAAGAGTA TTG<u>CA</u>T TAAAGTCTAACCTATAG GA<u>AT</u>CT TACAGCC<u>A</u>TCGAGAGGGACACGGCGAA), P* (TTATCAAAAAGAGTA TTG<u>T</u>CT TAAAGTCTAACCTATAG GAT<u>T</u>CT TACAGCC<u>A</u>TCGAGAGGGACACGGCGAA), and P** (TTATCAAAAAGAGTA TTG<u>TAA</u> TAAAGTCTAACCTATAG GAT<u>TT</u>T TACAGCC<u>A</u>TCGAGAGGGACACGGCGAA). Underlines indicate mutations from the original T7 A1 promoter. Ribosome binding site R is the RBS of gene 10 in the T7 phage (TTTAAGAAGGAGATATACAT),^37^ and R* (ATCACTCTACGGCCAGCTGCAAAC) was computationally designed using De Novo DNA version 2.1 to have a 10x weaker translation strength (14.8 A.U.) compared to R (148 A.U.).^38,54^ Primers used to change each of these elements are included in Table S1.

Plasmids and strains from this study and their sequences are available on AddGene (https://www.addgene.org/Mary_Dunlop/).

### Blue light exposure

Strains were grown overnight from a single colony in selective LB medium containing 100 μg/mL carbenicillin, 30 μg/mL kanamycin, or 25 μg/mL chloramphenicol as required for plasmid maintenance. Cultures were refreshed 1:100 in selective M9 minimal media (M9 salts supplemented with 2 mM MgSO_4_, 0.1 mM CaCl_2_, and 0.1% glucose) for two hours with 100 μM IPTG for induction of OptoCreVvd2 split recombinase production. Cultures were then either exposed to blue light or kept in the dark for two hours. Light exposure was performed using a light plate apparatus (LPA)^55^ with two 465 nM wavelength LEDs per well (ThorLabs LED465E), with an total output of 120 μW/cm^2^ per well.

### Minimum inhibitory concentration measurement

Minimum inhibitory concentrations (MICs) were measured based on the protocol outlined in Wiegand *et al*.^56^ Antibiotic stocks were made by dissolving the antibiotic in sterile distilled water (carbenicillin, kanamycin, tetracycline) or 99% ethanol (chloramphenicol), with concentrations normalized for potency based on CLSI standards.^57^ Assay plates for measuring the MIC were prepared by performing serial dilutions of antibiotic in 100 μL M9 minimal media in 96-well plates. Antibiotic concentrations were selected to include values that spanned the MIC levels for the dark and light state cultures in each experiment. Immediately following light exposure, cultures were normalized by dilution to the lowest optical density (OD) of each experiment. Normalized cultures were then diluted 1/25 into 96-well plates in triplicate and grown overnight for 18 hours at 37°C. The OD absorbance reading of each well was then measured at 600 nm (OD600) using a BioTek Synergy H1 plate reader.

### Colony forming unit measurement

Colony forming units (CFU) were measured following the micro-spotting protocol outlined in Sieuwerts *et al*.^58^ After MIC plates were grown overnight for 18 hours, cultures were serially diluted 1:10 in 1x M9 salts in a 96-well plate. Dilutions from 10^−1^ to 10^−6^ were spot-plated, with 5 μL on LB-agar plates in duplicate for each well (n = 6 for each condition). Plates were grown overnight at 37°C and colonies were counted by hand the next day for the lowest dilution with countable colonies for each sample. CFU counts per mL were then calculated by multiplying dilution level by the average number of colonies counted by per condition, normalizing for the 5 μL volume plated.

### Microscopy and image analysis

Strains were grown overnight from a single colony in selective LB medium. Cultures were refreshed 1:100 in selective M9 minimal media for two hours with 100 μM IPTG for induction of OptoCreVvd2. Samples were then placed on 1.5% low melting agarose pads made with M9 minimal media containing 100 μM IPTG and 400 μg/mL kanamycin (*knt* activation) or 60 μg/mL chloramphenicol (*cat* activation). Samples were grown at 30°C to prevent pads from drying out, and imaged every 15 minutes (*knt* activation) or 20 minutes (*cat*_*T172A*_ activation) for 18 hours. Cells were imaged at 100x using a Nikon Ti-E microscope. Blue light exposure was provided by a LED ring (Adafruit NeoPixel 1586), which was fixed above microscope stage and controlled by an Arduino with a custom Matlab script. Images were segmented and analyzed using the DeLTA 2.0 software and custom analysis scripts.^59^

### Statistical analysis

OD600 values in MIC curves and bar plots are reported as the mean of three samples ± the standard deviation. Colony count values for CFU measurements are reported as the mean of six samples consisting of two dilutions and platings for each MIC data point.

## Acknowledgements

We thank Heidi Klumpe for her helpful comments on the manuscript. We thank Caroline Blassick for creating the cat_T172A_ mutant. This work was supported by NIH grant R01AI102922 (MJD). MBS received support through the NIH training grant T32 EB006359.

## Author Contributions

MBS and MJD conceived and designed the experiments. MBS performed the experiments and analyzed the data. MBS and MJD wrote the manuscript.

## Competing Interests

The authors declare no competing financial interest.

## Notes

### Competing Interest Statement

The authors have declared no competing interest.

