## Supplementary Information for "An Optogenetic Toolkit for Light-Inducible Antibiotic Resistance"

**Supplementary Figures**

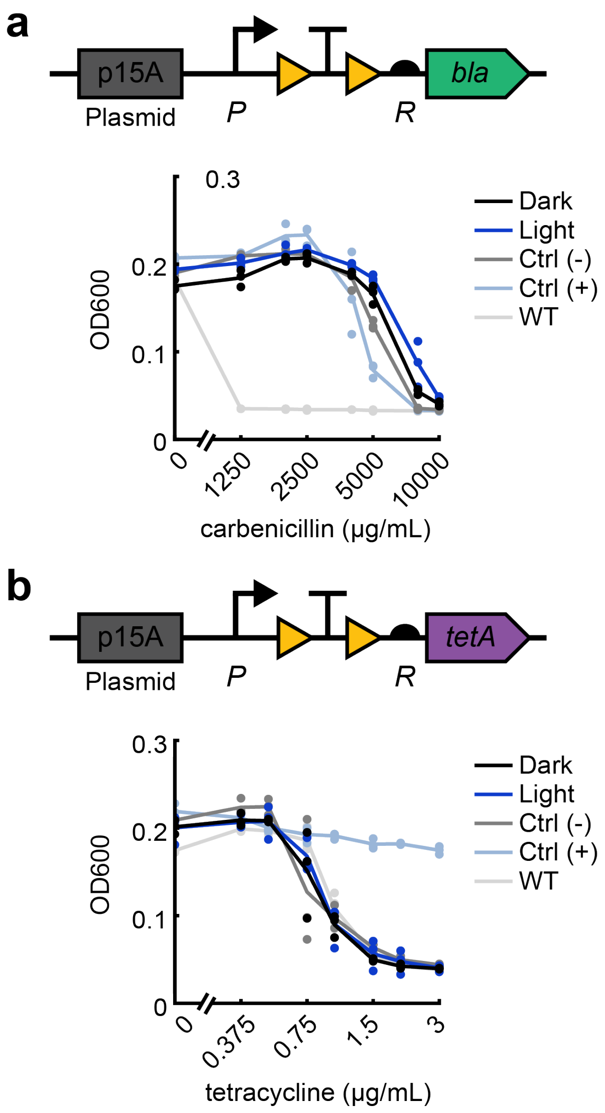


**Figure S1.** Optogenetic activation of **(a)** OptoCre-*bla* and **(b)** OptoCre-*tetA* on the p15A plasmid origin using promoter *P* and RBS *R*. MIC is quantified by OD600 after 18 hours (n = 3).

### Supplementary Movie Captions

**Movie S1.** Time-lapse microscopy of light-induced chromosomal OptoCre-*knt* with promoter P* and RBS R on agarose pads containing 400 μg/mL kanamycin (scale bar = 10 μm). Images show a representative position of the OptoCre-*knt* activation strain without (left) or with (right) blue light. Light is provided by an LED light ring above the microscope stage, and exposure begins immediately after cells are added to antibiotic-containing pads.

**Movie S2.** Time-lapse microscopy of light-induced p15A plasmid-based OptoCre-*cat* using $cat_{T172A}$ with promoter P and RBS R on agarose pads containing 60 μg/mL chloramphenicol (scale bar = 10 μm). Images show a representative position of the OptoCre-*cat* activation strain without (left) or with (right) blue light. Light is provided by an LED light ring above the microscope stage, and exposure begins immediately after cells are added to antibiotic-containing pads.

### Supplementary Tables

**Table S1.** Primers used for plasmid insertion of antibiotic resistance constructs using either Gibson or Golden Gate (GG) method. Binding regions in uppercase.

| Element | Assembly Method | Forward Primer | Reverse Primer |
|---|---|---|---|
| *bla* | Gibson | aagaaggagatatacatATGAGTATTCAACATTTCCG | tgccctggagatccctattaCCAATGCTTAATCAGTGAG |
| *knt* | Gibson | tttaagaaggagatatacatATGATTGAACAAGATGGATTGC | atgcctggagatccctattaTCAGAAGAACTCGTCAAGAAG |
| *cat* | Gibson | tttaagaaggagatatacatATGGAGAAAAAAATCACTGGAT | atgcctggagatccctattaTTACGCCCCGCCC |
| *tetA* (loxP-TT-loxP insertion) | GG | gtgactcgtctcgGCACGGCGAAATAACTTC | gtgactcgtctcgcgCTTCTTAAAATAACTTCGTATAATG |
| P* | GG | tacgctggtctccttGTCTTAAAGTCTAACCTATAGGATTCTTAC | tacgctggtctccgtCCCTCTCGATGGCTGTAAGA |
| P** | GG | tacgctggtctccttGTAATAAAGTCTAACCTATAGGATTTTTAC | tacgctggtctccgtCCCTCTCGATGGCTGTAAAA |
| R* | GG | gcatgaggtctcctaGCATACATTATACGAAGTTATATCACTCTACGG | cgatcaggtctcgaaTCATGTTTGCAGCTGGCCG |
| Plasmid resistance marker swaps | Gibson | GATCTATCAACAGGAGTCCAAGC | GCGCAACGCAATTAATGTAAGT |
| FRT cassette addition (cassette) | Gibson | gttttgcgccattcgatggtGCAGCATTACACGTCTTGAG | acatcaccgatggggaagatcctgtcaaacatgagaattaattccg |
| FRT cassette addition (vector) | Gibson | ttaattctcatgtttgacagGATCTTCCCCATCGGTGATGTC | ctcaagacgtgtaatgctgCACCATCGAATGGCGCAAAAC |
